# Tuftsin: a natural molecule against SARS-CoV-2 infection

**DOI:** 10.1101/2022.01.10.475746

**Authors:** Jiahao Huang, Jing Wang, Ziyuan Wang, Ming Chu, Yuedan Wang

## Abstract

Coronavirus disease 2019 (COVID-19) continuously proceeds despite the application of a variety of vaccines. It is still urgent to find effective ways to treat COVID-19. Recent studies indicate that NRP1, an important receptor of the natural peptide tuftsin, facilitates SARS-CoV-2 infection. Importantly, tuftsin is a natural human molecule released from IgG. Here, we found 91 overlapping genes between tuftsin targets and COVID-19-associated genes. Bioinformatics analyses indicated that tuftsin could also target ACE2 and exert some immune-related functions to treat COVID-19. Using surface plasmon resonance (SPR) analysis, we confirmed that tuftsin can bind ACE2 and NRP1 directly. Moreover, tuftsin effectively impairs the binding of SARS-CoV-2 S1 to ACE2. Thus, tuftsin is an attractive drug against COVID-19. And tuftsin as natural immunostimulating peptide in human, we speculate that tuftsin may has crucial roles in asymptomatic carriers or mild cases of COVID-19.

## Introduction

Coronavirus disease 2019 (COVID-19) caused by severe acute respiratory syndrome coronavirus 2 (SARS-CoV-2) results in high morbidity and mortality ^1,2^. It is known that the spike (S) protein binding to angiotensin-converting enzyme 2 (ACE2) is the core mechanism of SARS-CoV-2 infecting host cells. Through persistent efforts, COVID-19 vaccines have been approved for human use in most countries. However, the quantity of neutralizing antibodies induced by vaccines still needs to be verified in humans. At present, another hopeful intervention is neutralizing monoclonal antibodies (mAbs) ^3^. Unfortunately, producing safe and effective mAbs is complicated, and the duration of effective protection remains to be determined ^4,5^. Moreover, continuous mutations of SARS-CoV-2 during the pandemic may lead to escape from antibody recognition and reduce the neutralizing activity of mAbs ^6^. Hence, discovering a broad-spectrum and effective method for treating COVIV-19 is urgent.

Recently, neuropilin 1 (NRP1) has been found to be a host factor for SARS-CoV-2 infection ^7^. It has been reported that NRP1 facilitates the entry of SARS-CoV-2 into cells in the presence of ACE2 ^8^. It is worth noting that NRP1 is an important receptor of tuftsin ^9,10^. Tuftsin, a natural phagocytosis-stimulating peptide, was found by Victor Najjar et al. in 1970 ^11^. Tuftsin is released from the Fc fragment of IgG by an endocarboxy-peptidase in the spleen and a leukokininase on the outer membrane of neutrophilic leukocytes ^11,12^. Furthermore, tuftsin is a tetrapeptide that consists of Thr-Lys-Pro-Arg, located at amino acid residues 289 to 292 of the heavy chain of IgG. Tuftsin has a broad spectrum of activities mainly associated with immune system functions and exerts effects on phagocytic cells, especially macrophages. These functions of Tuftsin briefly include cell phagocytosis, motility, immunogenic response, and bactericidal and tumoricidal activities ^13,14^. It was reported that tuftsin activity is inversely correlated with splenectomy function and is significantly lower in patients with AIDS, cirrhosis, intestinal failure and some infectious diseases ^12,15,16^. Moreover, it was demonstrated that tuftsin has stability and low toxicity in vitro and in vivo ^14,17,18^. As a natural immune stimulating peptide, tuftsin is an attractive candidate for immunotherapy. Thus, we hypothesized that tuftsin could inhibit SARS-CoV-2 infection by interacting with NRP1. We subsequently performed experiments to verify our conceptions.

## Materials and Methods

### Compound profiling and disease-related gene identification

The structure of tuftsin was found in PubChem (https://pubchem.ncbi.nlm.nih.gov/). The 3D structure of tuftsin was built using Chem3D. Afterward, the target proteins corresponding to tuftsin screened from the Pharmmapper database and PubMed database were standardized in UniProt (http://www.uniprot.org/). Finally, Cytoscape 3.8.2 was used to determine the drug-target network. COVID-19-related genes were mined from the GeneCards database. All of the disease gene targets were normalized with R software using the Bioconductor package when redundancy was deleted ^19^.

### Network establishment

Screening for drug-disease crossover genes was performed. Based on previous steps, two sets of target lists were prepared: drug targets and disease-related genes. The crossover genes were filtered with R software using the Venn Diagram package. The STRING 11.5 database (http://string-db.org/) was used to analyse the intersecting protein–protein interactions (PPIs), and the common targets were counted with R software.

### Enrichment analysis

The proteins with overlapping expression patterns were evaluated by bioinformatics annotation with R software using the Bioconductor package, including a panther classification system (http://www.pantherdb.org/), a gene ontology (GO) annotation database website (http://www.geneontology.org), and Kyoto Encyclopedia of Genes and Genomes (KEGG) pathway enrichment analysis (http://www.genome.jp/kegg/). A p < 0.05 was considered statistically significant.

### Molecular docking analysis

The flexible docking process between tuftsin and target proteins was conducted by softwere Discovery Studio 2021 (DS). Briefly, the crystallographic structures of human ACE2 (PDB ID: 1R42) and human NRP1 (PDB ID: 2QQ1) with high resolution were prepared using the Prepare Protein and Minimization module of DS. The active binding site of each protein was defined based on the most representative features of the SARS-CoV-2 interface. Tuftsin was docked into the active binding site of ACE2 and NRP1 using the molecular docking module in DTS.

### Surface plasmon resonance analysis

The recombinant human ACE2 protein (Novoprotein, Beijing, China) and recombinant human NRP1 protein were used for surface plasmon resonance (SPR) analysis using a Biacore 8K instrument (Biacore, Uppsala, Sweden). Each target was immobilized onto flow cells in a CM5 sensor chip (GE Healthcare) via the amine-coupling method. Briefly, ACE2 and NRP1 were diluted in 10 mM pH 4.5 acetate to 20 μg/mL. Then, the protein solutions were injected individually on the carboxyl-modified sensor surface to form amine bonds. Both ACE2 and NRP1 immobilized levels were approximately 10000 RU. Binding analyses were carried out at 25°C and a flow rate of 10 μl/min. Tuftsin diluted in running buffer (1×PBS, 0.05% Tween 20 and 5% dimethyl sulfoxide, pH 7.4) was run over each target at gradient concentrations. An empty flow cell without any immobilized protein was used as a deducted reference. The binding curves were analysed using a kinetic binding model supplied with Biacore Evaluation Software (GE Healthcare).

### Competition binding experiment

For the competition binding experiment, the SARS-CoV-2 S1 protein was immobilized on the CM5 sensor chip via the amine-coupling method. 5 nM ACE2 was injected for negative control. Tuftsin was diluted into a series of solutions with gradient concentrations and fixed with 5 nM ACE2, and then the solutions were injected into the chip. The blocking efficacy was evaluated by comparison of response units with and without tuftsin incubation.

### Statistical analysis

The results were analysed using Student’s *t* test with SPSS software and R 4.1.0.

## Results

### Bioinformatics analyses revealed the connection between tuftsin and COVID-19

The 2D structure of tuftsin was obtained from the PubChem database (Compound CID: 156080), and the most stable 3D structure was built based on the 2D structure through a molecular simulation assay (Fig. 1A). In addition to the reported receptors of tuftsin, the potential targets of tuftsin in humans were also predicted through the PharmMapper database. Together, 284 targets of tuftsin were collected (Fig. 1B and data S1). Furthermore, we collected 2572 disease-associated genes of COVID-19 from the GeneCards database (data S2). We surprisingly found 91 intersecting proteins of tuftsin targets and COVID-19-associated genes through intersection analysis (Fig. 1C). It is intriguing that the overlapping proteins account for nearly one-third of tuftsin targets. Moreover, the protein–protein interaction network of the overlapping proteins was established, and it showed that JAK2, STAT1 and AKT1 are core molecules in the network (Fig. 1D). Furthermore, we performed enrichment analysis for the 91 intersecting genes. GO annotation revealed that the expressed tuftsin-COVID-19 crossover proteins were mainly associated with immune functions such as neutrophil activation, neutrophil-mediated immunity and cytokine receptor binding. Moreover, the KEGG pathway enrichment analysis showed that the COVID-19 pathway was the most significantly enriched. In addition, many target genes were strongly associated with some immunologic pathways, such as Th17 cell differentiation, the IL-17 signaling pathway and the immune checkpoint pathway (Fig. 1E). In the COVID-19 pathway, the SARS-CoV-2 receptors ACE2 and NRP1 were targets of tuftsin. Moreover, IL-2, STAT1 and some complement molecules in the COVID-19 pathway were targets of tuftsin (Fig. 1F). Together, these results suggest that tuftsin is a promising candidate against COVID-19, owing to its multifaceted pharmacological activities.

**Fig. 1.**
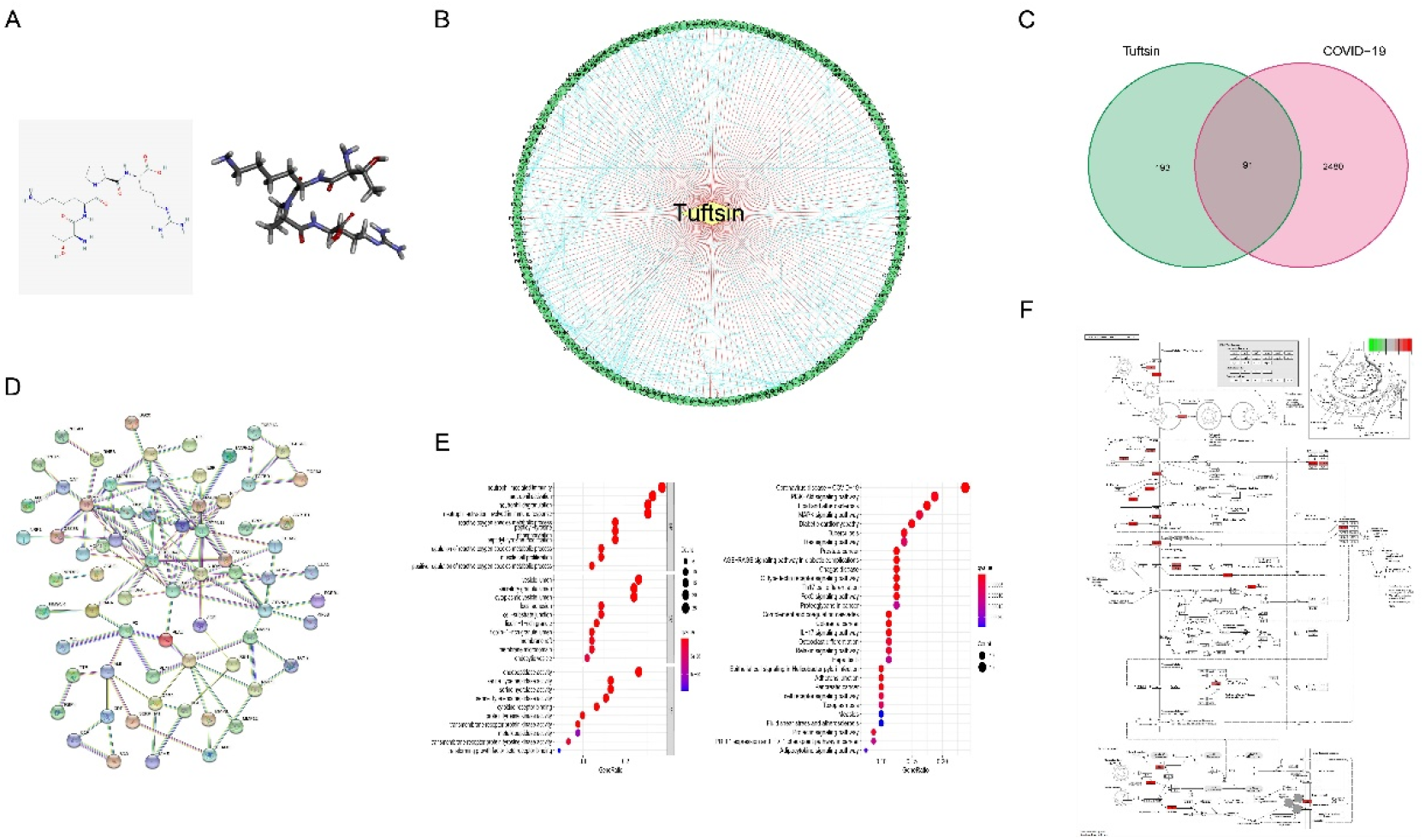
The connection between tuftsin and COVID-19. (**A**) (Left) The 2D chemical structure of tuftsin downloaded from the PubChem database. (Right) The 3D chemical structure of tuftsin established by software based on the 2D structure. (**B**) The ‘drug-target’ network of tuftsin. Red links represent the interactions between tuftsin and target nodes. Each node is a protein target. Green points represent the targeted proteins in humans. Blue links represent the interactions between the targets. (**C**) A Venn diagram of tuftsin and COVID-19 cotargeted genes. (**D**) Protein–protein interaction (PPI) network of the intersected targets. The interactions with a high confidence of 0.95. (**E**) (Left) Gene ontology enrichment results in bubble plot. (Right) The KEGG enrichment results in bubble plot. (**F**) Detailed targets of tuftsin in the COVID-19 pathway. Red points represent the tuftsin targets. The intensity of the color represents the possibility of tuftsin targeting. Deeper color indicates higher possibility.

### The interaction of tuftsin with ACE2 and NRP1 analysed by molecular docking

It is novel that ACE2 is a potential target of tuftsin, as mentioned above. Thus, molecular docking was performed to determine the potential binding sites and binding affinity between tuftsin and the SARS-CoV-2 receptors ACE2 and NRP1. First, we defined the interaction interface of SARS-CoV-2 S1-RBD with ACE2 as the active sites of ACE2. These interface sites in ACE2 include Q24, M82, N330, and R393, which are mainly located in the N-terminal peptidase domain of ACE2 ^20^. Then, the docking region was a sphere containing the defined ACE2 active sites (Fig. S1A). The results showed that the affinity of tuftsin and ACE2 was −6.9 kcal/mol, demonstrating that they could combine spontaneously (Fig. 2A). Furthermore, tuftsin could form strong hydrogen bonds to Ser47 and Asp67, carbon hydrogen bonds to His345, Asp67 and Asn51, and salt bridges to Asp67 of ACE2 (Fig. 2A). It is worth mentioning that the binding sites were adjacent to the interactional sites of S1-RBD and ACE2, indicating that tuftsin could inhibit S1 binding to ACE2 by covering their binding sites. Meanwhile, the b1b2 domain of NRP1 was prepared, as previous studies showed that the extracellular b1b2 domain of NRP1 mediates binding to CendR peptides ^21^. Then, the active sites of NRP1 b1b2 were defined according to the interactional sites of S1-RBD and NRP1 b1b2, including D320, E348, Y353 and so on ^7^. The docking region was a sphere containing the defined NRP1 b1b2 active sites (Fig. S1B). The docking results showed that tuftsin and NRP1 b1b2 have a high binding affinity of −8.1 kcal/mol. In addition, tuftsin solidly fits into a binding pocket on NRP1 b1b2 (Fig. 2B). Furthermore, tuftsin could form a salt bridge to Lys 397 and a carbon hydrogen bond to Pro398, which are near the interactional sites of S1-RBD and NRP1 b1b2. Moreover, the binding region of tuftsin and NRP1 overlapped with the binding area of NRP1 and S1-RBD in space (Fig. 2B). Collectively, these results demonstrated that tuftsin could bind ACE2 and NRP1 and inhibit the SARS-CoV-2 S1 binding of ACE2 and NRP1 by covering their interactional sites.

**Fig. 2.**
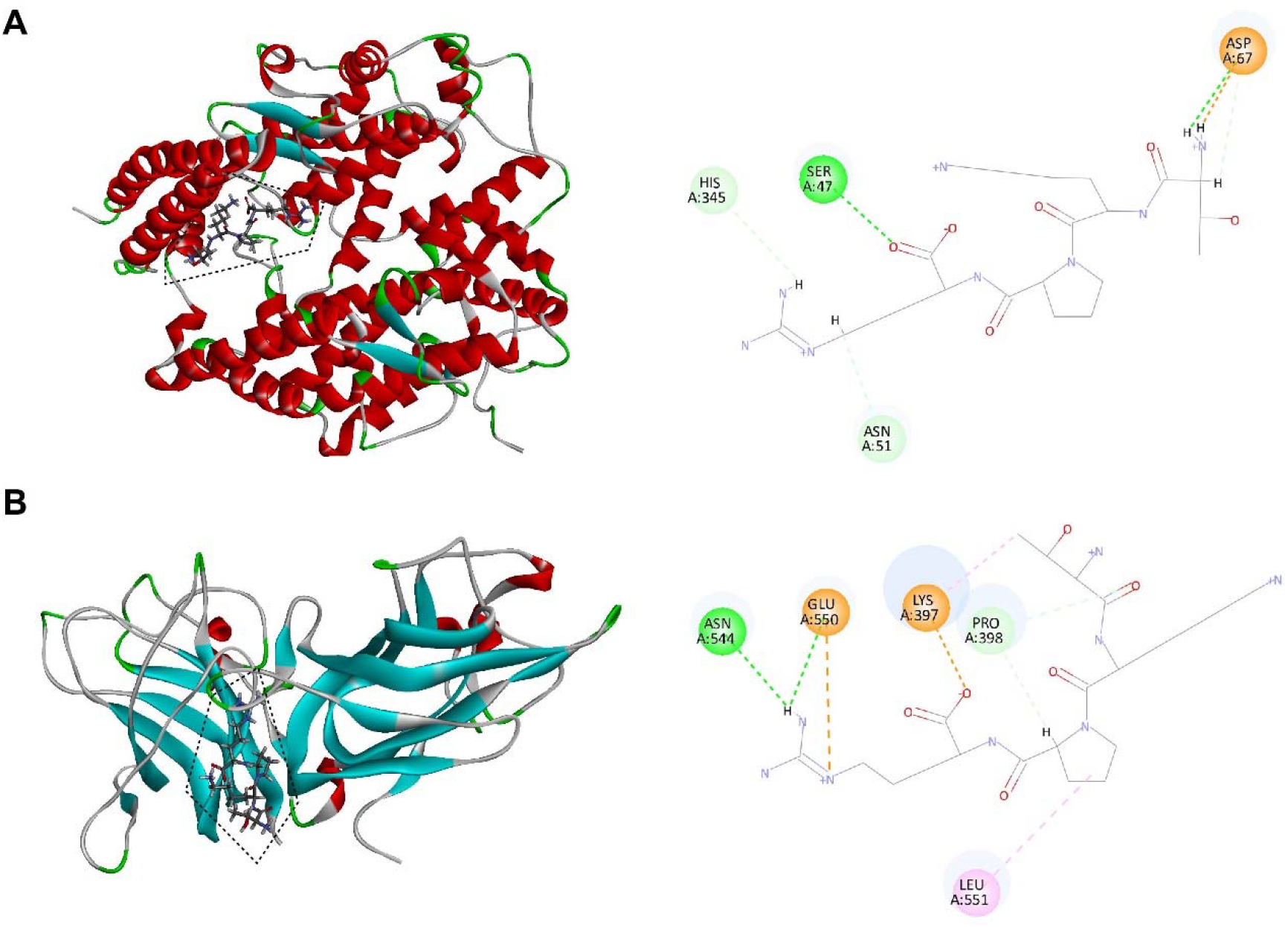
Molecular interaction of tuftsin with ACE2 and NRP1. (**A**) (Left) The binding pattern of tuftsin with ACE2. Binding area was circled by black dotted line. Secondary structural elements are depicted as ribbons (coils, α-helices, arrows, β-sheets). Color is based on secondary structures (α-helices, red; β-sheets, skyblue; loops, green). (Right) Molecular interaction schemes of tuftsin with the relative residues of ACE2. Green lines represent conventional hydrogen bonds; light green lines represent carbon hydrogen bonds; orange lines represent salt bridges; and pink lines represent alkyl bonds. (**B**) (Left) The binding pattern of tuftsin with NRP1. Binding area was circled by black dotted line. (Right) Molecular interaction schemes of tuftsin with the relative residues of NRP1. Other interpretations are the same as above.

### Tuftsin binds ACE2 and NPR1 directly, as confirmed by surface plasmon resonance (SPR) analyses

The interactions of tuftsin with ACE2 and NRP1 were further evaluated by real-time biomolecular interaction analysis with SPR. The kinetics of the binding reaction were determined by injecting different concentrations of tuftsin over recombinant human ACE2 immobilized on one half of the chip surface and over recombinant human NRP1 immobilized on another half of the chip surface. The results showed that tuftsin can bind ACE2 with an equilibrium dissociation constant (*K*D) of 460 μmol/L, according to the obtained association and dissociation rates (Fig. 3A). Moreover, the *K*D fitting curves of tuftsin and ACE2 became gentle with higher concentrations of tuftsin, indicating that the interaction of tuftsin and ACE2 is specific. (Fig. 3A). Tuftsin can also bind NRP1 specifically with a higher binding affinity of *K*D = 10.65 μmol/L. The sensorgrams and *K*D fitting curves of tuftsin and NRP1 are shown in Fig. 3B. As SPR is the gold standard for detecting drug-target interactions, these results demonstrate that tuftsin binds ACE2 and NRP1 directly and specifically, validating the accuracy of the above results of bioinformatics analyses and molecular docking assays.

**Fig. 3.**
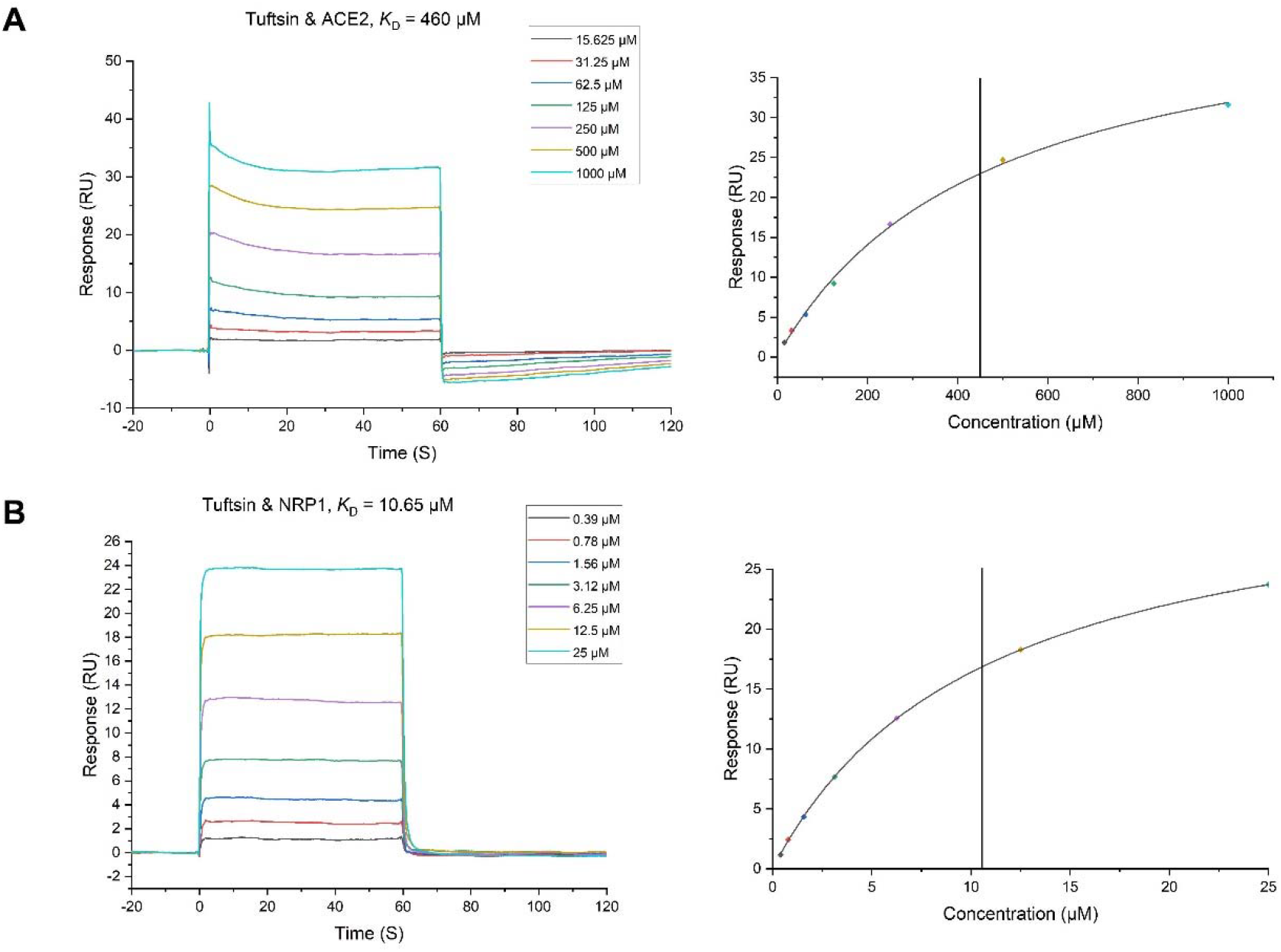
The binding of tuftsin to ACE2 and NRP1 was determined by SPR assay. (**A**) (Left) Binding curves of tuftsin with ACE2. The *K*D of the ACE2 protein with a series of concentrations of tuftsin was calculated by using a 1:1 binding model. Data are presented as response units (RU) over time (S). (Right) The fitting carve of tuftsin with ACE2. (**B**) (Left) Binding curves of tuftsin with NRP1. The *K*D of the NRP1 protein with a series of concentrations of tuftsin was calculated by using a 1:1 binding model. Other interpretations are the same as above. (Right) The fitting carve of tuftsin with NRP1.

### Tuftsin impairs the binding of SARS-CoV-2 S1 to ACE2

An SPR-based competition assay was employed to determine whether tuftsin could affect the binding of S1 protein with ACE2. We first determined the binding affinity of the S1 protein with ACE2 by SPR assay, which unsurprisingly showed a high affinity. A suitable concentration ACE2 solution was injected over the immobilized SARS-CoV-2 S1 protein as a control. Then, a series of gradient concentrations of tuftsin solutions containing equal concentrations of ACE2 were injected over the immobilized SARS-CoV-2 S1 protein for comparison. We observed that 9 μmol/L tuftsin had a mild inhibitory effect. It is worth noting that the addition of 156 μmol/L tuftsin significantly attenuated the response signal by approximately two-thirds compared to that of ACE2 alone over the immobilized S1. Notably, a substantial decrease in the response signal was observed with increasing concentrations of tuftsin. The response single was close to zero when the added concentration of tuftsin was 625 μmol/L. This result indicates that the interaction between S1 and ACE2 was almost completely blocked in the presence of 625 μmol/L tuftsin (Fig. 4). The experiment was repeated three times independently. In conclusion, the competition binding experiment revealed that tuftsin effectively impairs the binding of SARS-CoV-2 S1 to ACE2 in a dose-dependent manner.

**Fig. 4.**
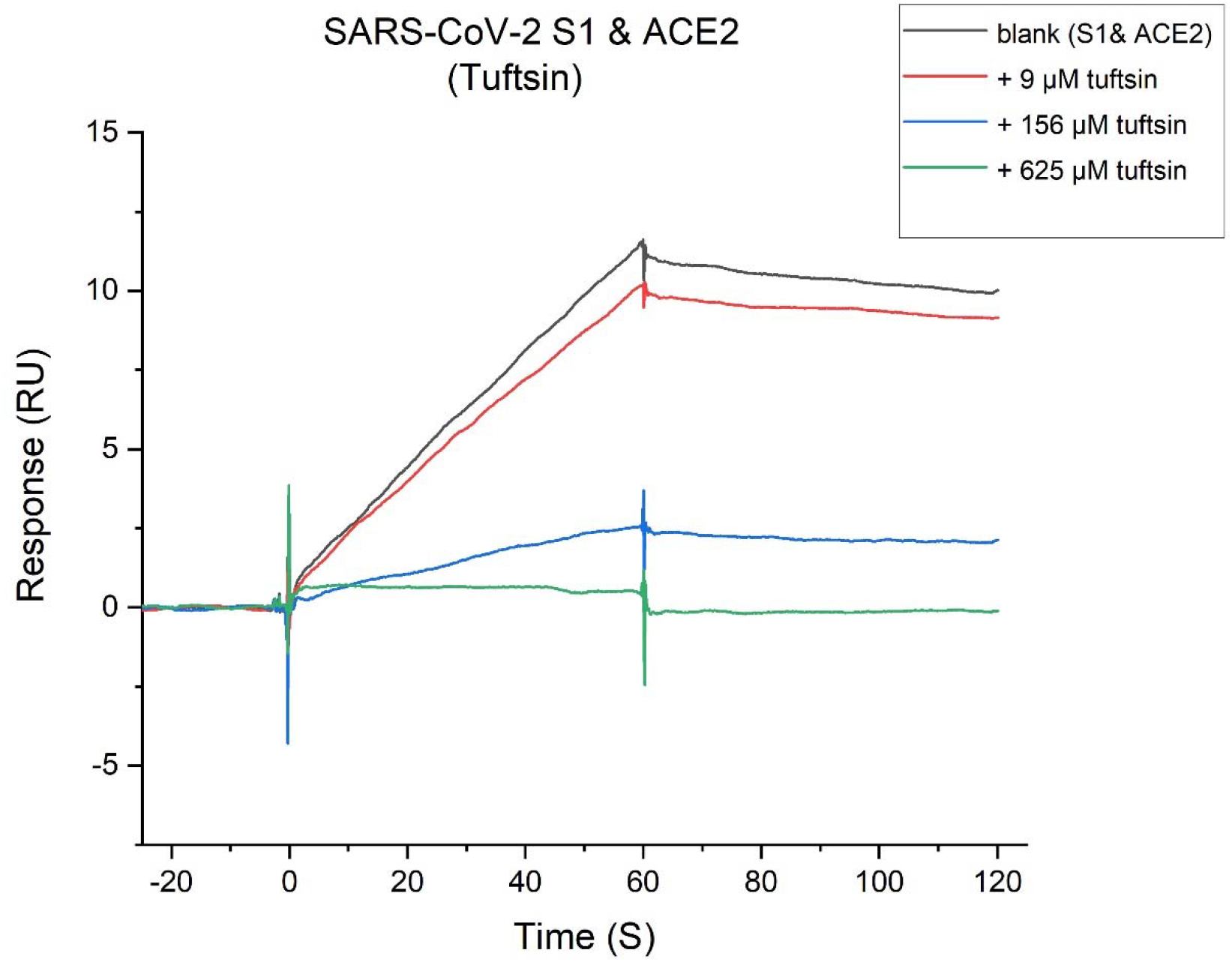
Tuftsin inhibits the SARS-CoV-2 S1 binding to ACE2. The binding activity of SARA-CoV-2 S1 to ACE2 in the presence of increasing concentrations of tuftsin. Intensive concentrations of tuftin showed enhanced inhibitory effects.

## Discussion

At present, vaccination is the most general way to prevent COVID-19; however, the notable problem is the uneven distribution of vaccine resources worldwide ^22^. The cost of producing vaccines and neutralizing antibodies is relatively high. It has been reported that the effectiveness of the SARS-CoV-2 vaccine declines significantly during 2021 ^23^. Here, we report that an immune-stimulating peptide, tuftsin, is a potential effective drug for COVID-19. Tuftsin, as a natural tetrapeptide that exists in humans, originates from a special fraction of the parent carrier IgG through enzymatic processing. Accordingly, tuftsin has lower toxicity and fewer side effects than other drugs ^24^. There are many marked drugs, such as oral liquid of spleen aminopeptide, which mainly contains tuftsin, and some drugs, which are derivatives of tuftsin, which all have satisfactory clinical efficacy ^25^. Importantly, tuftsin can be produced on a large scale at a lower cost ^26^. This allows tuftsin to be widely applied for the prevention and treatment of COVID-19 infection. The general existence of tuftsin in species allows wide protection of animals. It is worth noting that the mutant sequence of tuftsin turns inactive or inhibitory analogs ^27^.

In this research, 9 μM tuftsin had slight inhibitory activity. We observed that when the concentration of tuftsin was 156 μM, the binding affinity of SARS-CoV-2 S1 and ACE2 was reduced significantly. When the concentration reached 625 μ M, the combination of SARS-CoV-2 S1 and ACE2 was completely blocked. It has been confirmed that a 156 μM concentration of tuftsin can exist at a high concentration in the internal environment after intravenous injection ^28^. We conceive that tuftsin can be designed as an oral or nasal spray. In this case, the local concentration of tuftsin reached 625 μM. It has been reported that the amount of IgG induced by vaccines is mainly focused on the lower respiratory tract. Consequently, the upper respiratory tract, which mainly suffers from viral infection, lacks sterilizing immune protection ^29^. Importantly, the spray form of tuftsin could protect the upper respiratory tract, which the antibodies induced by vaccines cannot effectively protect. The molecular docking assay showed that tuftsin binds at the N-terminus of ACE2, which is the area of S1 protein binding. This indicated that tuftsin can block the binding of S1 protein and ACE2 directly.

It is worth noting that there were many asymptomatic and mild infectors during the pandemic. It is clear that innate and adaptive immunity functions during asymptomatic infection; however, the mechanism of the T cell and antibody response is unclear ^30,31^.

Asymptomatic people seem to clear the virus faster ^32^. Tuftsin, a human natural immunostimulating peptide released from IgG, certainly has significant roles related to innate immunity. We reported that tuftsin can target the important receptors of SARS-CoV-2 S1, which is similar to adaptive immunity. Thus, we speculated that tuftsin has crucial roles in asymptomatic or mild infection. It is likely that the activity of tuftsin is higher in asymptomatic individuals than in symptomatic individuals.

## Conflict of Interest

The authors declare that the research was conducted in the absence of any commercial or financial relationships that could be construed as a potential conflict of interest.

## Author contributions

Y.W., J.H. and C.M. conceptualized and designed this study. J.H. performed the bioinformatic analysis. J.H. and Z.W. performed the molecular docking. J.H. performed the SPR experiments with assistance from J.W. J.H. processed the data. J.H. drafted, edited the manuscript. Y.W., C.M. administered the project. All authors have read and agreed to the published version of this manuscript.

## Funding

National Natural Science Foundation of China (81603119) and Natural Science Foundation of Beijing Municipality (7174316).

## Acknowledgements

We thank the State Key Laboratory of Natural and Biomimetic Drugs (Peking University) for their assistance in performing the surface plasmon resonance assay.

## Supplementary Material

Figs. S1

Data S1 and S2

## Data Availability Statement

All data are available in the main text or the supplementary information.

## Notes

### Competing Interest Statement

The authors have declared no competing interest.

## References and Notes

1 Zhu, N. et al. A Novel Coronavirus from Patients with Pneumonia in China, 2019. N Engl J Med 382, 727–733, doi:10.1056/NEJMoa2001017 (2020).

2 Dai, L. & Gao, G. F. Viral targets for vaccines against COVID-19. Nat Rev Immunol 21, 73–82, doi:10.1038/s41577-020-00480-0 (2021).

3 Renn, A., Fu, Y., Hu, X., Hall, M. D. & Simeonov, A. Fruitful Neutralizing Antibody Pipeline Brings Hope To Defeat SARS-Cov-2. Trends Pharmacol Sci 41, 815–829, doi:10.1016/j.tips.2020.07.004 (2020).

4 Su, W. et al. Neutralizing Monoclonal Antibodies That Target the Spike Receptor Binding Domain Confer Fc Receptor-Independent Protection against SARS-CoV-2 Infection in Syrian Hamsters. mBio 12, e0239521, doi:10.1128/mBio.02395-21 (2021).

5 Taylor, P. C. et al. Neutralizing monoclonal antibodies for treatment of COVID-19. Nat Rev Immunol 21, 382–393, doi:10.1038/s41577-021-00542-x (2021).

6 Du, L., Yang, Y. & Zhang, X. Neutralizing antibodies for the prevention and treatment of COVID-19. Cell Mol Immunol 18, 2293–2306, doi:10.1038/s41423-021-00752-2 (2021).

7 Daly, J. L. et al. Neuropilin-1 is a host factor for SARS-CoV-2 infection. Science 370, 861–865, doi:10.1126/science.abd3072 (2020).

8 Cantuti-Castelvetri, L. et al. Neuropilin-1 facilitates SARS-CoV-2 cell entry and infectivity. Science 370, 856–860, doi:10.1126/science.abd2985 (2020).

9 Vander Kooi, C. W. et al. Structural basis for ligand and heparin binding to neuropilin B domains. Proc Natl Acad Sci U S A 104, 6152–6157, doi:10.1073/pnas.0700043104 (2007).

10 von Wronski, M. A. et al. Tuftsin binds neuropilin-1 through a sequence similar to that encoded by exon 8 of vascular endothelial growth factor. J Biol Chem 281, 5702–5710, doi:10.1074/jbc.M511941200 (2006).

11 Najjar, V. A. & Nishioka, K. “Tuftsin”: a natural phagocytosis stimulating peptide. Nature 228, 672–673, doi:10.1038/228672a0 (1970).

12 Corazza, G. R. et al. Tuftsin deficiency in AIDS. Lancet 337, 12–13, doi:10.1016/0140-6736(91)93331-3 (1991).

13 Najjar, V. A. Tuftsin, a natural activator of phagocyte cells: an overview. Ann N Y Acad Sci 419, 1–11, doi:10.1111/j.1749-6632.1983.tb37086.x (1983).

14 Fridkin, M. & Najjar, V. A. Tuftsin: its chemistry, biology, and clinical potential. Crit Rev Biochem Mol Biol 24, 1–40, doi:10.3109/10409238909082550 (1989).

15 Zoli, G. et al. Impaired splenic function and tuftsin deficiency in patients with intestinal failure on long term intravenous nutrition. Gut 43, 759–762, doi:10.1136/gut.43.6.759 (1998).

16 Trevisani, F. et al. Impaired tuftsin activity in cirrhosis: relationship with splenic function and clinical outcome. Gut 50, 707–712, doi:10.1136/gut.50.5.707 (2002).

17 Siemion, I. Z. & Kluczyk, A. Tuftsin: on the 30-year anniversary of Victor Najjar’s discovery. Peptides 20, 645–674, doi:10.1016/s0196-9781(99)00019-4 (1999).

18 Amoscato, A. A., Davies, P. J., Babcock, G. F. & Nishioka, K. Receptor-mediated internalization of tuftsin by human polymorphonuclear leukocytes. J Reticuloendothel Soc 34, 53–67 (1983).

19 Yu, G., Wang, L. G., Han, Y. & He, Q. Y. clusterProfiler: an R package for comparing biological themes among gene clusters. Omics 16, 284–287, doi:10.1089/omi.2011.0118 (2012).

20 Lan, J. et al. Structure of the SARS-CoV-2 spike receptor-binding domain bound to the ACE2 receptor. Nature 581, 215–220, doi:10.1038/s41586-020-2180-5 (2020).

21 Plein, A., Fantin, A. & Ruhrberg, C. Neuropilin regulation of angiogenesis, arteriogenesis, and vascular permeability. Microcirculation 21, 315–323, doi:10.1111/micc.12124 (2014).

22 Forni, G. & Mantovani, A. COVID-19 vaccines: where we stand and challenges ahead. Cell Death Differ 28, 626–639, doi:10.1038/s41418-020-00720-9 (2021).

23 Cohn, B. A., Cirillo, P. M., Murphy, C. C., Krigbaum, N. Y. & Wallace, A. W. SARS-CoV-2 vaccine protection and deaths among US veterans during 2021. Science 0, eabm0620, doi:doi:10.1126/science.abm0620.

24 Catane, R. et al. Toxicology and antitumor activity of tuftsin. Ann N Y Acad Sci 419, 251–260, doi:10.1111/j.1749-6632.1983.tb37111.x (1983).

25 Shakya, N., Sane, S. A., Haq, W. & Gupta, S. Augmentation of antileishmanial efficacy of miltefosine in combination with tuftsin against experimental visceral leishmaniasis. Parasitol Res 111, 563–570, doi:10.1007/s00436-012-2868-z (2012).

26 Siebert, A., Gensicka-Kowalewska, M., Cholewinski, G. & Dzierzbicka, K. Tuftsin - Properties and Analogs. Curr Med Chem 24, 3711–3727, doi:10.2174/0929867324666170725140826 (2017).

27 Najjar, V. A. Biochemical aspects of tuftsin deficiency syndrome. Med Biol 59, 134–138 (1981).

28 Blok-Perkowska, D., Muzalewski, F. & Konopińska, D. Antibacterial properties of tuftsin and its analogs. Antimicrob Agents Chemother 25, 134–136, doi:10.1128/aac.25.1.134 (1984).

29 Krammer, F. SARS-CoV-2 vaccines in development. Nature 586, 516–527, doi:10.1038/s41586-020-2798-3 (2020).

30 Boyton, R. J. & Altmann, D. M. The immunology of asymptomatic SARS-CoV-2 infection: what are the key questions? Nat Rev Immunol, 1–7, doi:10.1038/s41577-021-00631-x (2021).

31 Schijns, V. & Lavelle, E. C. Prevention and treatment of COVID-19 disease by controlled modulation of innate immunity. Eur J Immunol 50, 932–938, doi:10.1002/eji.202048693 (2020).

32 Nogrady, B. What the data say about asymptomatic COVID infections. Nature 587, 534–535, doi:10.1038/d41586-020-03141-3 (2020).

