## Supplementary Material for "Tuftsin: a natural molecule against SARS-CoV-2 infection"

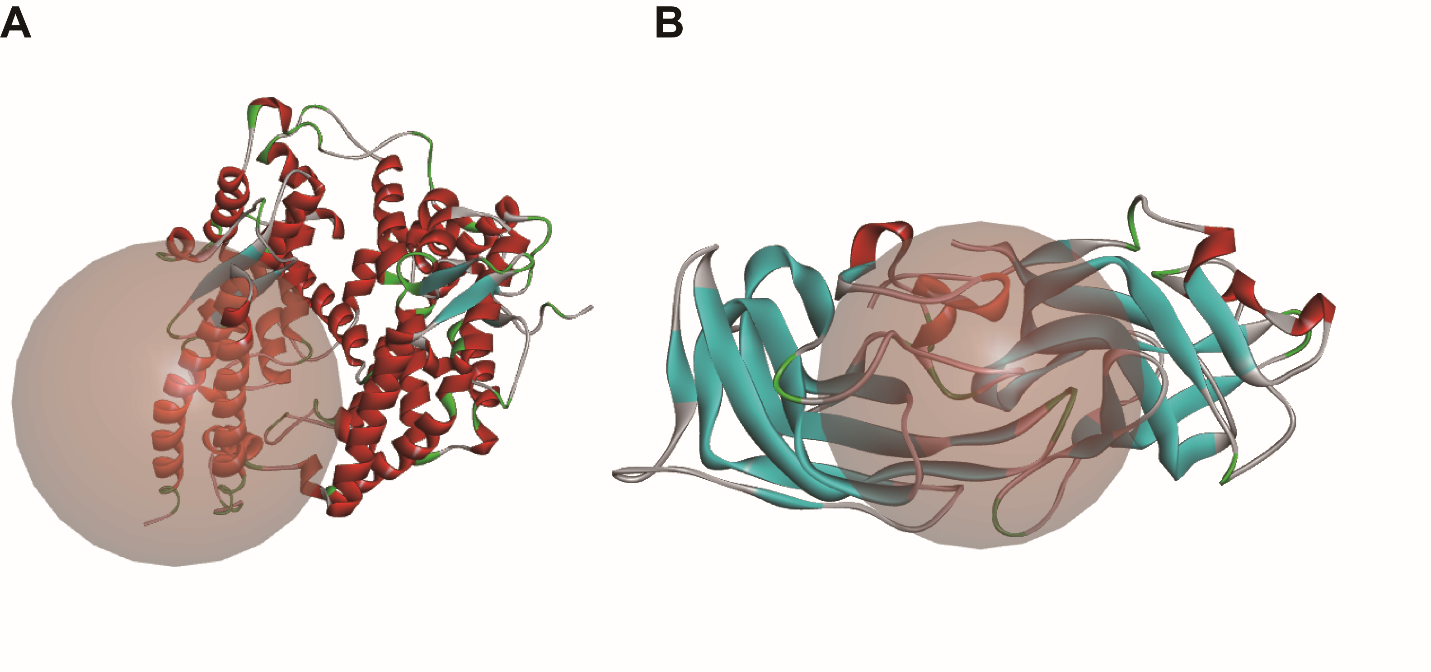

Fig. S1.

(A) The defined binding region of ACE2. The sphere represents the defined binding region. The center sites of the sphere were based on the interfaced sites of ACE2 and SARS-CoV-2 S1. (B) The defined binding region of NRP1. The sphere represents the defined binding region. The center sites of the sphere were based on the interfaced sites of NRP1 and SARS-CoV-2 S1.

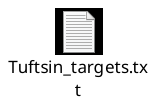

Data S1.

The 284 tuftsin targets were collected from published and predicted results.

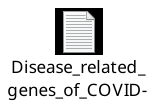

**Data S2.**

The 2572 disease-related genes of COVID-19 were collected from GeneCards.
